# The intertwined metabolism of *Medicago truncatula* and its nitrogen fixing symbiont *Sinorhizobium meliloti* elucidated by genome-scale metabolic models

**DOI:** 10.1101/067348

**Authors:** Thomas Pfau, Nils Christian, Shyam K. Masakapalli, Lee J. Sweetlove, Mark G. Poolman, Oliver Ebenhöh

## Abstract

Genome-scale metabolic network models can be used for various analyses including the prediction of metabolic responses to changes in the environment. Legumes are well known for their rhizobial symbiosis that introduces nitrogen into the global nutrient cycle. Here, we describe a fully compartmentalised, mass and charge-balanced, genome-scale model of the clover *Medicago truncatula*, which has been adopted as a model organism for legumes. We employed flux balance analysis to demonstrate that the network is capable of producing biomass (amino acids, nucleotides, lipids, cell wall) in experimentally observed proportions, during day and night. By connecting the plant model to a model of its rhizobial symbiont, *Sinorhizobium meliloti*, we were able to investigate the effects of the symbiosis on metabolic fluxes and plant growth and could demonstrate how oxygen availability influences metabolic exchanges between plant and symbiont, thus elucidating potential benefits of amino acid cycling. We thus provide a modelling framework, in which the interlinked metabolism of plants and nodules can be studied from a theoretical perspective.

## 1 Introduction

Nitrogen belongs to the elements which are absolutely crucial for life, because it is contained in all amino acids and many other essential biomolecules. Whereas nitrogen is the most abundant element in the earth’s atmosphere, most of it is present in the inert form of nitrogen gas (N_2_) constituting approximately 80% of the total atmosphere. Therefore, despite its high abundance, nitrogen is often a limiting factor for growth in plants. To overcome this, in agriculture large amounts of nitrogen are applied to the soil in the form of artificial fertilisers to promote plant growth. Before the industrial revolution, farmers applied a crop rotation scheme to their fields and left the field barren or planted it with beans or peas every few years to let the soil ‘recover’. We now know that this strategy re-introduces bio-available nitrogen into the soil. This is due to ability of the planted crops (beans or peas) or plants quickly occupying barren fields, such as clovers, to form a nitrogen fixing symbiosis with rhizobia. Rhizobia carry a gene for the highly oxygen sensitive nitrogenase enzyme, which catalyses the reduction of atmospheric nitrogen to ammonium, and the plants provide the symbiont with an environment protecting it from damaging oxygen. In this way atmospheric nitrogen is introduced into the global organic nutrient cycle. This symbiosis has seen the most attention and is best understood in plants of the Fabacaea (or Leguminosae) family, to which beans, peas and clovers belong (Brewin, 1991; Ferguson et al., 2010; Hellriegel et al., 1888; Hirsch et al., 2001; Drevon et al., 2015).

In recent years, *Medicago truncatula* has become a model plant for the legume-rhizobia symbiosis (Cook, 1999). Low nitrogen availability will trigger the recruitment of rhizobia to the plant roots and initiate the nodulation, in which rhizobia invade the plant root and root nodules are formed. In these nodules, the rhizobia are taken up by plant cells, surrounded by a membrane and differentiate into bacteroids. Upon completing differentiation they begin fixing nitrogen, which is made available to the plant primarily in the reduced form of ammonium. In return, the plant provides organic acids as nutrients to the rhizobia (Ferguson et al., 2010; Udvardi and Poole, 2013). In addition there is evidence that amino acid cycling is essential for nitrogen fixation at least in some rhizobial strains (Lodwig et al., 2003; Prell et al., 2010). Obviously, the mutual dependence of the metabolism of plant and rhizobium is crucial for this symbiotic interaction. During the establishment of the nodules, the metabolic fluxes in both plant and rhizobium change drastically (Prell and Poole, 2006).

To explore, understand and analyse the changes in the metabolic fluxes during nodulation, computer simulations are a highly useful tool, which becomes increasingly established in modern biology research. For this, detailed error-free metabolic network models have to be established. Whereas there exist now over a hundred (Baart and Martens, 2012; Monk et al., 2014) genome-scale metabolic network models for a wide range of organisms, and a number of steps in the development of these models can now efficiently be automated (Dias et al., 2015; Caspi et al., 2016), the construction of genome-scale metabolic network models still involves numerous manual curation steps and therefore is very time-consuming (Fell et al., 2010). However, even before model construction is finished and established simulation techniques, such as flux balance analysis (FBA) can be employed, the model building process itself provides considerable insight into the metabolic capabilities of the investigated organism and allows to refine its genome annotation. Once a highly curated metabolic network model is established, it presents a theoretical framework allowing to query the system and understand its functional properties. The numerous possible theoretical investigations (Rezola et al., 2015) are for example useful to help understanding the structure and regulation of metabolic networks (Poolman et al., 2009; Nikerel et al., 2012), identify essential genes (Joyce and Palsson, 2008), predict putative drug targets (Perumal et al., 2011) or support engineering of novel pathways (Basler et al., 2012) producing desired compounds of technological or economic interest. In the plant sciences, genome-scale models with different degrees of accuracy and completeness exist for the model unicellular green alga *Chlamydomonas reinhardtii* (Chang et al., 2011), the model species *Arabidopsis thaliana* (Mintz-Oron et al., 2012; Poolman et al., 2009), rice *Oryza sativa* (Poolman et al., 2013) and others (de Oliveira Dal’Molin et al., 2010; Saha et al., 2011). While initially plant models were unicellular representations of metabolism, not distinguishing between different tissues, recent efforts tend to refine these models and generate multi-tissue representations (Gomes De Oliveira Dal’molin et al., 2015). In this paper, we focus on the presentation of a highly curated genome-scale metabolic model of *M. truncatula* and provide an analysis of general metabolic properties of its biochemical reaction network. For the construction of our detailed and curated model, which contains 1636 reactions in 7 compartments, we have developed a novel approach to assign metabolic reactions to sub-cellular compartments. Our approach integrates a range of available experimental data with computational predictions to construct a compartmentalised metabolic network model from an uncompartmentalised model. We further progressed the quality of our curation by ensuring that all reactions and transport processes are not only mass-balanced, but also charge-balanced. This is an important improvement over other genome-scale network models because it allows for a system wide prediction of fluxes of protons and other charged particles over cellular membranes, which was restricted to small subsystems in earlier models (Poolman et al., 2009). Charge balancing over membranes is a key prerequisite to realistically describe electron transfer chains required for ATP biosynthesis. Compartmentalisation allows for a more precise distinction between the function and necessity of isozymes which are present in multiple compartments. Omitting this information would allow the model to use reactions for which substrates are not available, because they are not transported between compartments. This would make the prediction of potentially lethal mutations less reliable. We further refined the model by integrating tissue specific information to better represent the distribution of metabolism to root and shoot of the plant.

We discuss insights gained from the reconstruction process itself and their consequences on the annotation of genes with previously unknown functions and the refinement of annotations of previously insufficiently annotated genes. We experimentally obtained tissue specific biomass data of M. *truncatula*, thus providing for the first time a system wide in-depth biomass composition for this plant. Our detailed model makes it possible to assess precise energy and reductant requirements for the production of all metabolites in the biomass and we identify compounds which are in principle producible during the night from starch without the need for respiration.

To investigate the interaction between plant and microbe, this highly curated plant model was connected to a model of *Sinorhizobium meliloti*, derived from the MetaCyc database (Caspi et al., 2016), and we investigated the effect of nitrogen fixation and symbiosis on the plant model. We further addressed the question which fluxes might be most restrictive for nitrogen fixation and investigated the effect of small alterations of oxygen supply on the symbiotic nitrogen fixation capacity.

The model presented in this paper provides an important resource for researchers in plant sciences in general and for studying plant-rhizobia interactions in particular. Our simulations help to interpret and understand responses of plant metabolism, which include rearrangements of metabolic fluxes and changes in energy and reductant requirements, as a result of changes in nitrogen availability.

## 2 Results

### Properties of the *Medicago truncatula* metabolic network

Starting from the annotated genome version 3.5v5 (Young et al., 2011) and the biochemical information in the MedicCyc (v1.0) (Urbanczyk-Wochniak and Sumner, 2007) and BioCyc (Caspi et al., 2010) databases, we first built a database using the PathwayTools program (Karp et al., 2010).

Based on this refined database, we next built a metabolic network model without information about intracellular compartments (see Methods). After an initial manual curation to fill gaps (see Gap Filling), this network contained 1636 reactions. To get an initial overview of the biochemical processes involved, we have grouped these reactions into the different categories of metabolism, based on the BioCyc categorisation. Figure 1 shows the numbers of reactions assigned to the different MetaCyc pathway categories. While a majority of reactions (1117) are involved in biosynthesis pathways, only 294 are involved in degradation processes. Several reactions are present in both categories, because some processes are involved in both biosynthesis and degradation pathways.

**Figure 1:**
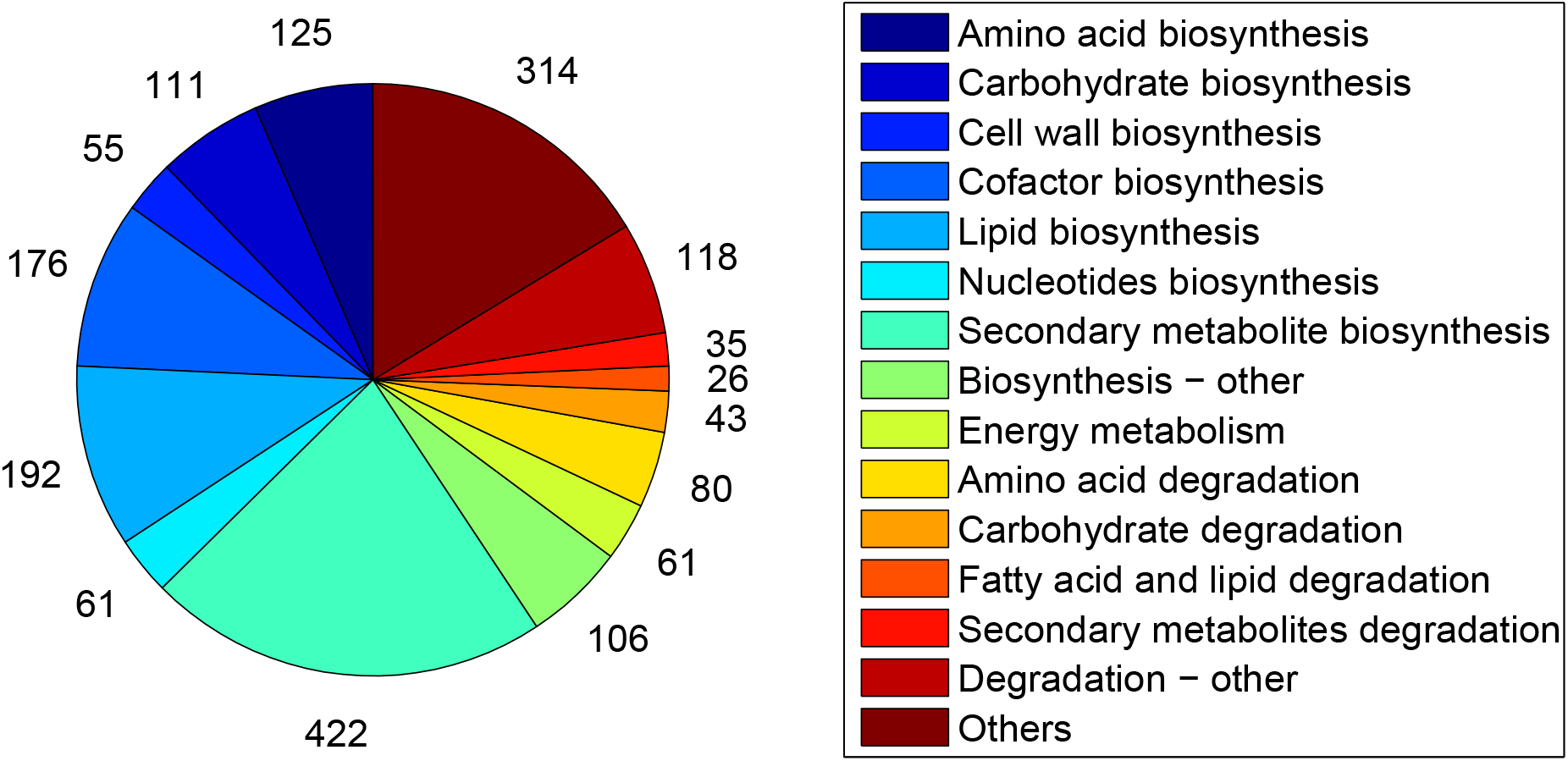
Categorisation of the reactions in the model based on MetaCyc pathway categories. 1117 reactions are involved in biosynthesis processes and 293 reactions are involved in degradation pathways, with 113 reactions being present in both categories. There are several biosynthetic reactions involved in multiple biosynthesis processes and a single degradation reaction involved in multiple degradation processes. The category ‘Others’ comprises all reactions present in the network, which are not transporters, importers or exporters and are either not assigned to any pathway or not included in any of the categories shown.

To fully compartmentalise our model and assign the 1636 reactions to the 7 compartments (cytosol, mitochondria, plastid, peroxisome, endoplasmic reticulum, vacuole and Golgi), we applied the procedure detailed in the Methods section. In short, first experimental evidence for sub-cellular localisation (Daher et al., 2010; Dubinin et al., 2011; Heazlewood et al., 2007) was used to assign the corresponding reactions to a compartment. Subsequently, the network extension method (Christian et al., 2009) was applied to add reactions to the different compartments, ensuring essential biochemical functions associated with each compartment (for details see Methods and Supplementary Material 1). All reactions that remained unlocalised after this step were then assigned to the cytosol. A final manual curation step assured that metabolites producible in the uncompartmentalised network were still producible in the compartmentalised network in at least one compartment. Figure 2 gives an overview of the compartmentalisation process and the assignment of the reactions to the different compartments. The large increase in the number of reactions associated with the ER in the final curation step results from a manual assignment of various sterol biosynthesis pathways to the ER (Benveniste, 2004).

**Figure 2:**
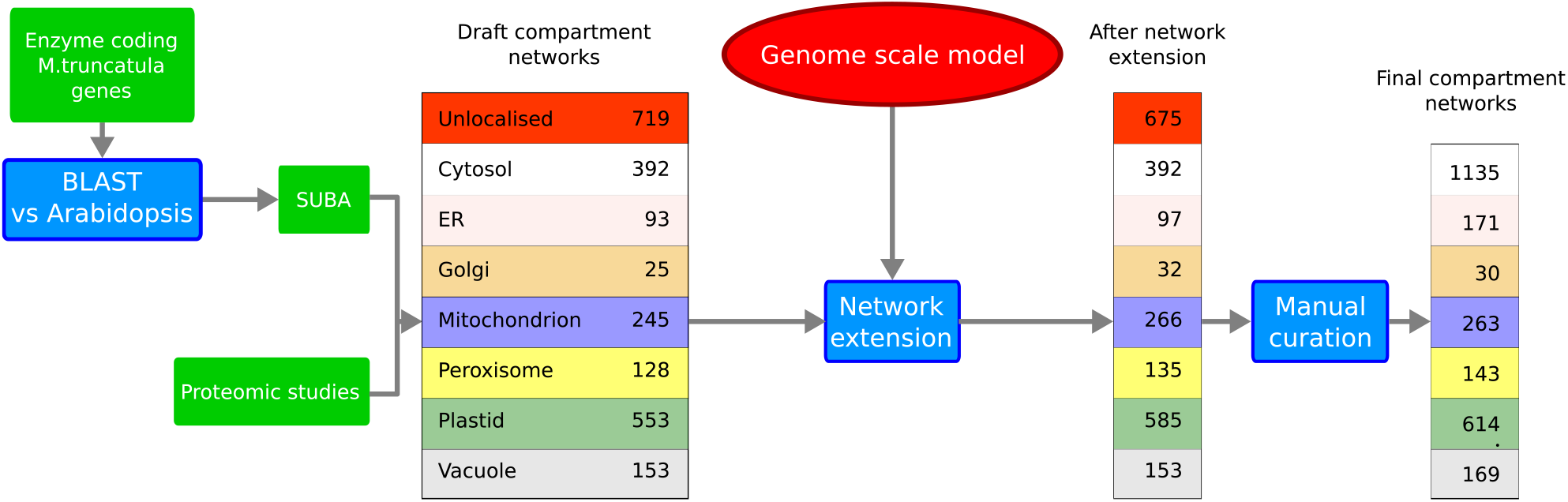
Process of compartment localisation. Assigning compartments to reactions was performed in three steps. The first step comprises a BLAST search for Arabidopsis homologues and obtaining proteomic data (Daher et al., 2010; Dubinin et al., 2011). In a second step, network extension (Christian et al., 2009) is applied to ensure that every organelle can perform key functions. The uncompartmentalised genome-scale model serves as a reference network. In a third step, it is ensured by manual curation that all compounds are producible which were producible in the uncompartmentalised network.

An overview of the metabolism is represented in Figure 3. Roughly a third (572) of the reactions are present in more than one compartment leading to a total of 2526 reactions in the model. For most of these reactions their multiple localisation was inferred based on homology to Arabidopsis (Heazlewood et al., 2007) or evidence from proteomic studies (Daher et al., 2010; Dubinin et al., 2011) while few were predicted by the network extension algorithm. The compartments are connected by a total of 271 transporters taken from the literature (Babujee et al., 2010; Helliwell et al., 2001; León and Sánchez-Serrano, 1999; Linka and Weber, 2010) or inferred in the final gap filling step (see Materials and Methods). The model reactions connect 1370 distinct chemical species, of which 701 are present in more than one compartment, resulting in a total of 2780 unique metabolic species in the model. Genetic evidence is present for over 70% of all non transport reactions.

**Figure 3:**
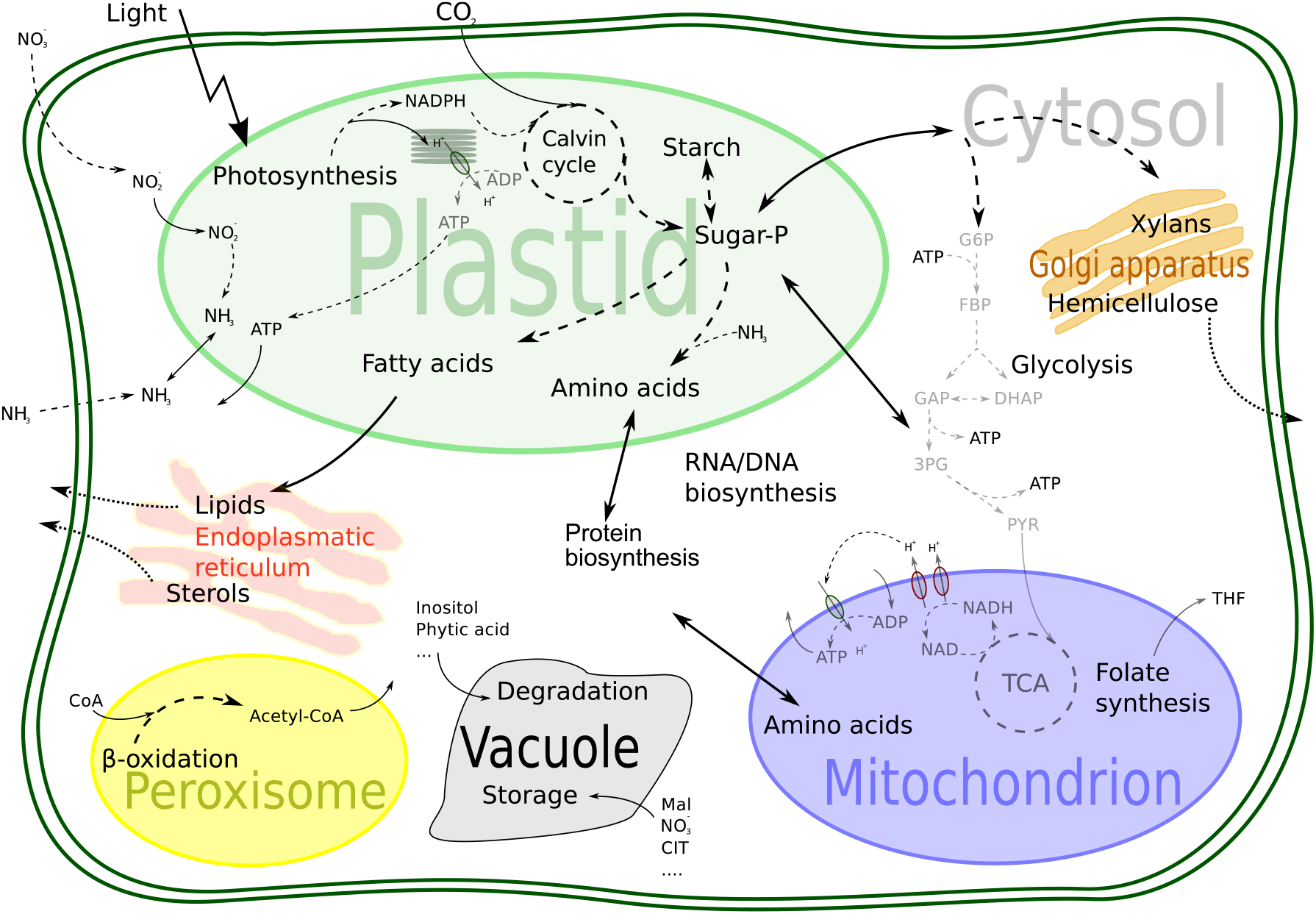
Schematic overview of the metabolic processes and compartments included in the genome-scale model of *M. truncatula*.

### Suggested re-annotation

While searching for enzymatic evidence for reactions filling gaps in the ascorbate biosynthesis pathway, we identified gene *MTR_4g092750* (Entrez Gene ID: 11406810), a homologue of galactose dehydrogenase *(AT4G33670* (Entrez Gene ID: 829509, BLAST E-Value = 1e–149, 78% sequence identity), that provides an enzymatic function which was initially lacking in the model. We therefore suggest a specification of the current annotation, which classifies this gene as a general aldo/keto reductase family oxidoreductase. During the biomass analysis of *M. truncatula* we could determine pinitol as present in the plant. When trying to determine the biosynthetic pathway for this substance we found that the enzyme encoded by gene *MTR-4g03844-0* (Entrez Gene ID: 11446905), currently annotated as a caffeic acid 3-O-methyltransferase, is very similar (87% sequence identity) to the inositol methyltransferase from *Glycine max* (Entrez Gene ID: 100812768). Since we could not find other candidates for this biosynthetic step, we expect that this enzyme might indeed be mis-annotated and would suggest a revision of its annotation.

### Experimental biomass composition and growth rates

For the computational analysis applied here, it is essential to know growth rates and the composition of the biomass. Biomass components were measured from extracts of eight week-old plants grown hydroponically. The major biomass fractions included protein, methanol soluble metabolites, lipids, starch, cell wall, chlorophyll and nucleotides (see Table II). Experiments were performed to obtain compositions for roots, stems and leaves. Based on Mettupalli (2011) we assumed a 1:1:1 distribution of leaf, stem and root material for the full plant composition. The composition of the protein (amino acids) and soluble extracts (sugars, free amino acids, organic acids etc) were measured by GC-MS. Chlorophyll was determined spectrometrically and total lipid content and cell wall content was measured gravimetrically. The composition of the latter fractions (cell wall, lipid) was taken from literature or assumed to be similar to data for *Medicago sativa*. Fatty acid composition data from Bakoglu et al. (2010) was used. Cell wall composition, including cellulose, pectins, lignin and hemicellulose, was derived from data of Jung and Engels (2002), Nakashima et al. (2008) and Johnson et al. (2007) and assumed to be the same for *M. truncatula* as for *M. sativa*.

**Table I:** Flux constraints for metabolite exchange between rhizobium and plant. For conversion between bacteroid and plant weight a ratio of 8.4mg/g was assumed.

| Flux | Experimental value<br>$nmol\ mg^{-1}\ min^{-1}$<br>(based on bacteroid weight) | Model constraint<br>$\mu mol\ g^{-1}\ h^{-1}$<br>(based on plant weight) | Source |
| --- | --- | --- | --- |
| Succinate uptake | 22.1 | 0 - 9.4248 | (1) |
| Malate uptake | 18.7 | 0 - 11.1384 | (1) |
| Total dicarboxylate uptake | - | 0-11.1384 |  |
| Amino acid uptake | 75.8 | 0 - 37.99* | (1) |
| Oxygen uptake | 14.1 - 39.2 | 16.8*<sup>2</sup> | (2) |

\*: This constraint was used to limit the total amino acid exchange (uptake and export).
\*<sup>2</sup>: The listed oxygen uptake is in the upper range of possible uptake rates. When alternate values are used this is explicitly mentioned.
(1):McRae et al. (1989) , (2):Bergersen (1997)

**Table II:**
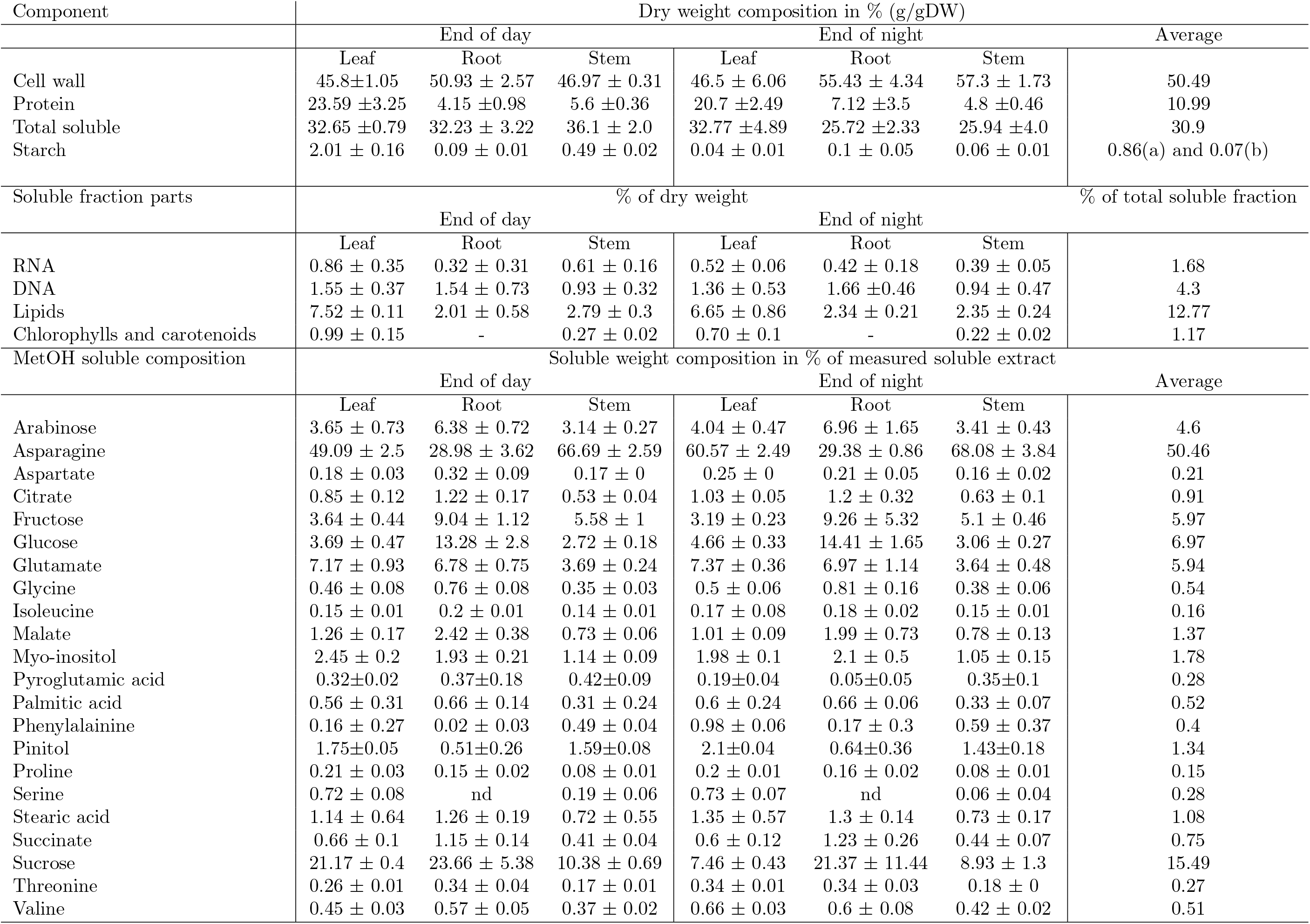
Biomass composition of *M. truncatula*. Measurements were taken after night period and after the light period. The soluble composition was determined using standard curves and obtaining percentages of the total measured amounts. For the biomass composition used in the analysis the average values were used and pyroglutamate was assumed to have spontaneously cyclized from glutamate. The MetOH soluble amount for the calculations is assumed to be the remaining fraction of the total soluble amount. Errors represent the standard deviation from three technical replicates, nd - not detected.

Starch content was determined at the end of day and end of night. The accumulation of starch was 19.62 ± 1.66 and 4.25 ± 0.27mgg^−1^ dry-weight per day for leaves and stems, respectively. Roots only showed minor changes in starch levels between end of day and end of night with 0.88 ± 0.02 and 0.98 ± 0.05 mgg^−1^ respectively, so we assumed these levels to be constant. The daily increase of starch in leaves and stem has to be put into the perspective of the overall growth of the plant. Taking common growth rates of plants into account – 0.1gg^−1^ d^−1^ to 0.12gg^−1^ d^−1^ for *M. sativa* (Lötscher et al., 2004) or 0.12gg^−1^ d^−1^ to 0.25gg^−1^ d^−1^ for *A. thaliana* (Pyl et al., 2012) – this means that the plant has to produce much higher levels of starch during the day than any other compound. Because of its special role as a storage compound, starch was not assumed to be part of the biomass: In contrast to other biomass compounds, which have to be produced in an amount reflecting the relative growth rate, starch has to be produced as a reserve for the night, regardless whether the plant is growing fast or slow. This is illustrated by a simple calculation based on our experimentally determined leaf starch content change of 19.62 mg g^−1^ dry-weight per day and an assumed daily growth rate between 12 to 25%. This results in a daily starch production in a leaf corresponding between 16.35% for 12% growth rate down to 7.85% for 25% growth rate of the total biomass and a subsequent conversion of a large fraction of this starch during the following night.

The experimentally determined biomass composition was used in our computational analyses to constrain the production of biomass to the observed composition. The ‘biomass reaction’ further contains the energy (ATP) demand required for the polymerisation of nucleic acids (DNA and RNA) and amino acids. It should be noted, however, that maintenance energy requirements are not included due to reasons discussed below.

### A multi-tissue model of *Medicago truncatula*

A (non tissue-specific) genome-scale model encompasses all biochemical reactions catalysed by enzymes, which are encoded in the genome. Thus, such a model can reflect the summed metabolic capabilities of an organism, but is unable to describe processes in a particular cell type or tissue. We therefore constructed a multi-tissue model of *Medicago truncatula* employing the FASTCORE (Vlassis et al., 2014) algorithm (see Materials and Methods for details). Here, we focus on describing two tissues only, namely root and shoot. However, in principle our approach can be generalised to more tissues. Based on the full genome-scale network, the employed FASTCORE algorithm generates smaller subnetworks, which represent the active metabolism in a particular tissue. This resulted in a reduced network size totalling 2151 reactions. The shoot tissue was represented by 1067 reactions (containing 168 internal transporters) and the root tissue by 1051 reactions (162 transporters). These submodels were connected by 32 inter-tissue transporters and a combined biomass reaction. The large reduction can partly be explained by many reactions in secondary metabolism which, in the current model formulation, are unable to carry flux, because specific exchange reactions have not been added. In total, the consistent part of the *Medicago truncatula* network contained only 1303 reactions, which are able to carry non-zero flux with the current configuration of exchange reactions, while all remaining reactions can be activated if additional exporters are included. The two-tissue model was able to sustain growth during day (using light as energy source and photosynthesis for carbon) and night (with starch as energy and carbon source). In addition, both ammonium and nitrate can be used as nitrogen sources to support growth under day and night conditions.

### Computational analysis

#### Network self-consistency

During cellular growth, all intermediate metabolites are diluted and have to be replenished. These metabolites are not included in the biomass, but the necessity to reproduce them results from the fact that cells grow and divide (Benyamini et al., 2010; Kruse and Ebenhöh, 2008). We therefore first tested whether the model is self-consistent in the sense that all metabolites can be produced from the nutrients using constraint-based modelling (see Methods). We found that 1161 of the 1370 metabolites could be produced. The remaining 209 metabolites fall into three main classes.

The first class, containing 121 metabolites, includes those substances which form conserved quantities. Examples are the acetyl carrier proteins linked to fatty acids, macromolecular redox carriers, such as ferredoxins, and tRNAs, which are necessary for the incorporation of amino acids into proteins. Most of these macromolecules are small proteins and RNA molecules which are implicitly contained in the biomass as part of the protein and nucleic acid fractions. Since the model can produce biomass (see below), biosynthesis is possible for over 86% of the compounds in this class. The remaining are small molecules, which cannot be biosynthesized because they are themselves needed for their own biosynthesis. An example is quinate, for which no biosynthetic route is present in MetaCyc, where it is only included as a substrate for degradation. However it is used in one of two routes from coumaroyl-CoA to caffeoyl-CoA, in which it is needed as a carrier compound.

The second class contains 65 metabolites, which are not connected to the remaining network. This group contains many of the biotin biosynthesis pathway intermediates (6 compounds). The reason why this pathway is disconnected is that it is unknown how the pimeloic acid moiety necessary for biotin biosynthesis is synthesised in plants. Most of the other metabolites (37) in this class are produced by specific enzymes, but no pathway for the production of their precursors is known.

The third class consists of 23 metabolites, which result from the degradation of modified macromolecules (e. g. polyfructofuranose), or which are present only in a degradation pathway for xenobiotics (e. g. cyanate). This phenomenon of metabolites which can only be ‘produced’ by degradation was discussed in detail earlier (Christian et al., 2009).

#### Biomass production in the genome scale model

We confirmed that the model is able to reflect fluxes in a living cell by maintaining production of biomass, with a composition as measured, from the mineral nutrients. We ensured that this is possible under two conditions: growth in light, where the light reactions provide energy and reductants, and growth in darkness, where carbon and energy is derived from transitory starch. For the simulations of the dark condition, the ability to utilise light to drive photosynthesis is turned off setting the respective flux to zero. For both conditions, sucrose and glucose importers are disabled. *In silico* knockout experiments predict that, with starch as carbon and energy source, 236 reactions are essential, among which are 26 transporters, including amino acid and fatty acid transporters from the plastid and cell wall precursor transporters to the Golgi apparatus. Under phototrophic growth, 243 reactions are essential, including 25 transporters. In total, 232 reactions are essential under both conditions. Starch degrading reactions are essential only under night conditions, while those involved in the Calvin cycle and photosynthesis are essential only in light conditions. Interestingly, the additional transporter predicted as essential under dark conditions is the maltose exporter in the chloroplast envelope (MEX). However, *mex* mutants are viable, albeit with a reduced growth and starch excess phenotype (Niittyl et al., 2004). The reason that the model predicts this transporter as essential is that, in the strict mathematical sense, no flux distribution exists which fulfils the stationarity condition for maltose. Therefore, the prediction can be understood in terms of the mathematical model formulation, and the predicted phenotype (maltose accumulation) is one of the key characteristics of the *mex* mutant. Of the essential metabolic reactions (excluding transporters) over 90% have genetic evidence. Our knockout simulations further revealed an additional set of 16 and 18 reactions for growth in the dark and light, respectively, which are essential in the sense that they must be present in at least one compartment, but in the compartmentalised model they have been assigned to several compartments. The two additional reactions for which at least one isozyme has to be present in light conditions are the triose isomerase reaction and phosphoglycerate kinase, which are both necessary to utilise Calvin cycle-derived triose phosphates. Using the objective to minimise the total flux through the system while producing a given amount of biomass (see Methods), we found that a set of 430 (dark) and 423 (light) reactions, of which 72 are transporters, are sufficient to produce all biomass components.

#### Energy requirements for biomass precursors

During night, plants use the stored starch as carbon and energy source. To investigate the nightly energy metabolism, it is interesting to know how much energy and reductant are minimally required to build essential building blocks, such as nucleotides, amino acids and organic acids. The theoretically predicted minimal energy and reductant requirements for selected compounds are depicted in Fig. 4, when using as nitrogen source either ammonium (clear bars) or nitrate (shaded bars). Obviously, the requirements depend critically on the nitrogen source. Growing on ammonium, most nucleotides and some amino acids, along with some organic acids and sugars can theoretically be directly produced from starch without additional ATP obtained through respiration (a full list including the reductant and energy costs is provided in the Supplementary Material 3). In fact, some of the compounds can be synthesised from starch while even producing a small surplus of energy equivalents. In contrast, with nitrate as nitrogen source, the synthesis of all amino acids and nucleotides requires additional ATP and NAD(P)H, for which some of the starch needs to be respired. This is not surprising because the reduction of nitrate to ammonia requires four reducing equivalents. Using nitrate exclusively, only organic acids are still producible from starch without additional energy demands. Because they do not contain nitrogen, their requirements are independent on the nitrogen source. Our calculations are in agreement with earlier observations that heterotrophic cell cultures of *A. thaliana* grown on nitrate tend to produce higher levels of organic acids and sugars and lower levels of amino acids than cells grown on mixed ammonium and nitrate (Masakapalli et al., 2013).

**Figure 4:**
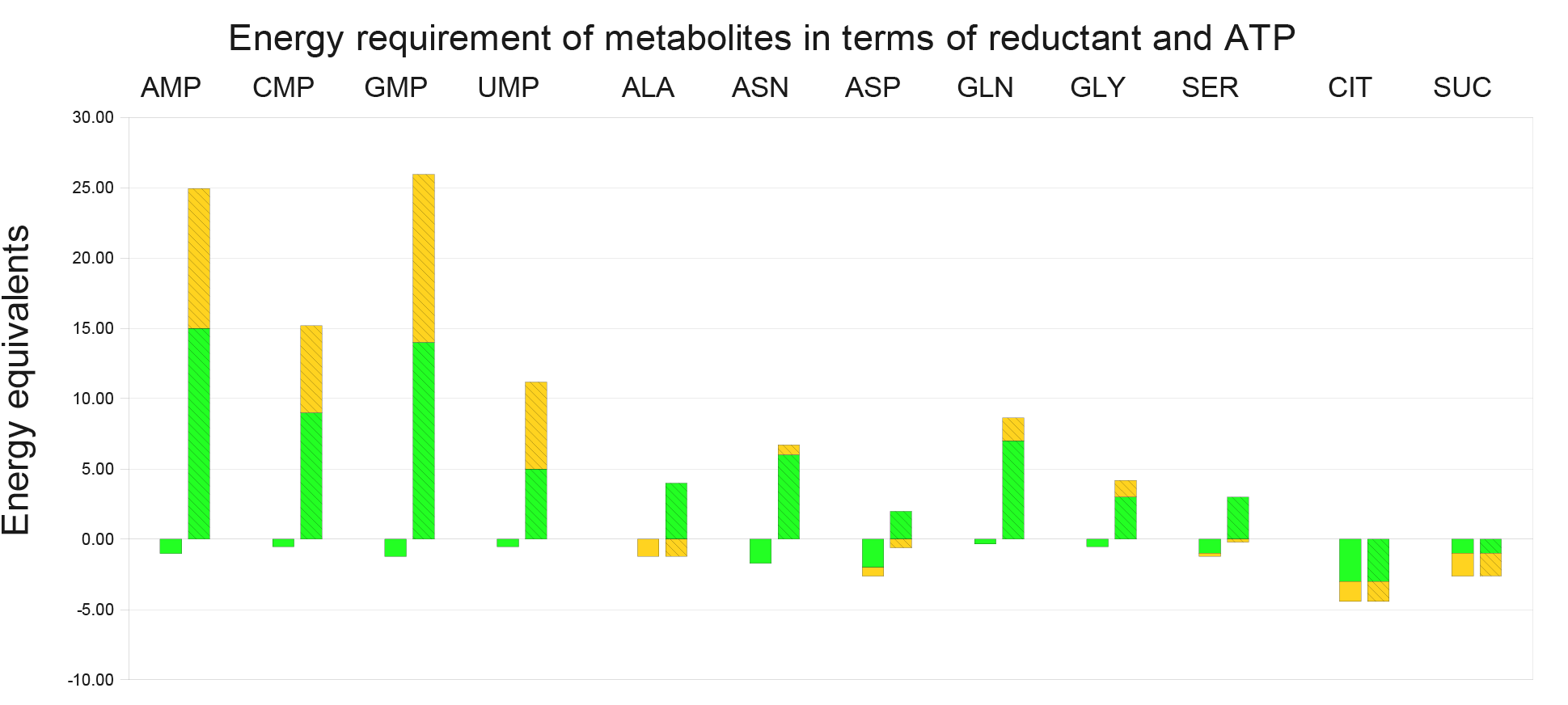
Theoretically calculated energy and reductant balance to produce important metabolites from starch. The bars indicate the amount of ATP (yellow) and NADPH (green) equivalents, which are minimally required to produce the respective metabolite (see Methods). Negative numbers depict a surplus. Numbers are shown for ammonium (clear) and nitrate (hatched) as the only nitrogen source.

These findings give rise to the question how much carbon contained in transient starch can *in principle* be used to produce biomass and how much needs to be respired in order to provide the necessary energy. To answer this question we calculated flux distributions producing a given amount of biomass under two objectives. The first objective demanded that the resulting flux distribution minimises the overall flux through all reactions (flux minimisation – FM) assuming equal weights for all reactions, and the second minimised the carbon which is exported (as CO_2_ or other organic compounds). The second objective gives the theoretical optimum of carbon conversion from starch to biomass.

Again, the results depend strongly on the nitrogen source. We found that, growing on ammonium, 92.2% of starch carbon can theoretically be converted into biomass (FM yielded the slightly lower value of 92.0%). On nitrate, however, a lower fraction of 79.2% (78.2% under FM) can theoretically be converted. In all cases, the remaining fraction of starch-derived carbon was respired and exported as CO_2_. Similarly as for the single compounds, the reduced fractions on nitrate can readily be explained by the fact that the reduction of nitrate requires a considerable amount of reductant. It is interesting that both objectives give very similar results. The small discrepancies are explained by the fact that FM favours shorter pathways, which sometimes are less energy efficient. It should be stressed that these figures overestimate the carbon yield, because the calculation neglects any energy or reductant requirement for maintenance purposes. Still, it is remarkable that our results reflect experimentally observed differences in growth rates on nitrate and ammonia (Brix et al., 2002) and comparison to real growth rates during the night (see Discussion) may be helpful to understand the quantity and nature of maintenance metabolism.

#### Nitrogen metabolism in the two tissue model

In the calculations above, we simulated the extreme scenarios that exclusively either nitrate or ammonium is available as nitrogen source during the night. To investigate the relevant intermediate cases, we simulated conditions in which both sources are available. However, supplying both nitrogen sources and directly applying any of the discussed objectives would lead to the trivial result that the model selects ammonium as the source, because it is ‘cheaper’ in terms of energy and reductant.

To simulate a transition, we therefore restricted the amount of nitrogen available to the plant and scanned over a range of different ammonium vs nitrate compositions. To calculate the fluxes, we fixed the growth rate to 0.1 g/gDW per day and minimised the sum of the squares of all fluxes in the system. In Fig. 5 selected flux responses are depicted, where the x-axis reflects the amount of available ammonium, which ranges from zero to the uptake rate required to sustain the growth rate of 0.1 g/gDW per day using only ammonium as nitrogen source. As expected, the amount of required starch (dash-dotted dark blue) drops with increasing availability of ammonium. This directly reflects the higher costs for nitrate assimilation. However, with more available ammonium, the activity of the mitochondrial respiratory chain and, concomitantly, the mitochondrial ATP synthase (dotted light blue) are increased. This observation can be explained by considering the changes in redox balance: More nitrogen available as ammonium results in a reduced reductant requirement to reduce nitrate. Therefore, more of the reductant produced in the TCA cycle (dotted green) can be used to generate ATP. The plot simultaneously shows the change of proton (dashed orange line) and carbonate (dashed purple) export to the soil. This uptake and release of protons and carbonate is mostly due to the required charge balance, since our model does not include inorganic ions. While a large fraction will, in nature, be balanced by the uptake of either positive or negative ions (Marschner, 1991), this computational results still nicely illustrates and at least partially explains the acidifying properties of ammonium nutrition in contrast to nitrate nutrition (Alam et al., 2007).

**Figure 5:**
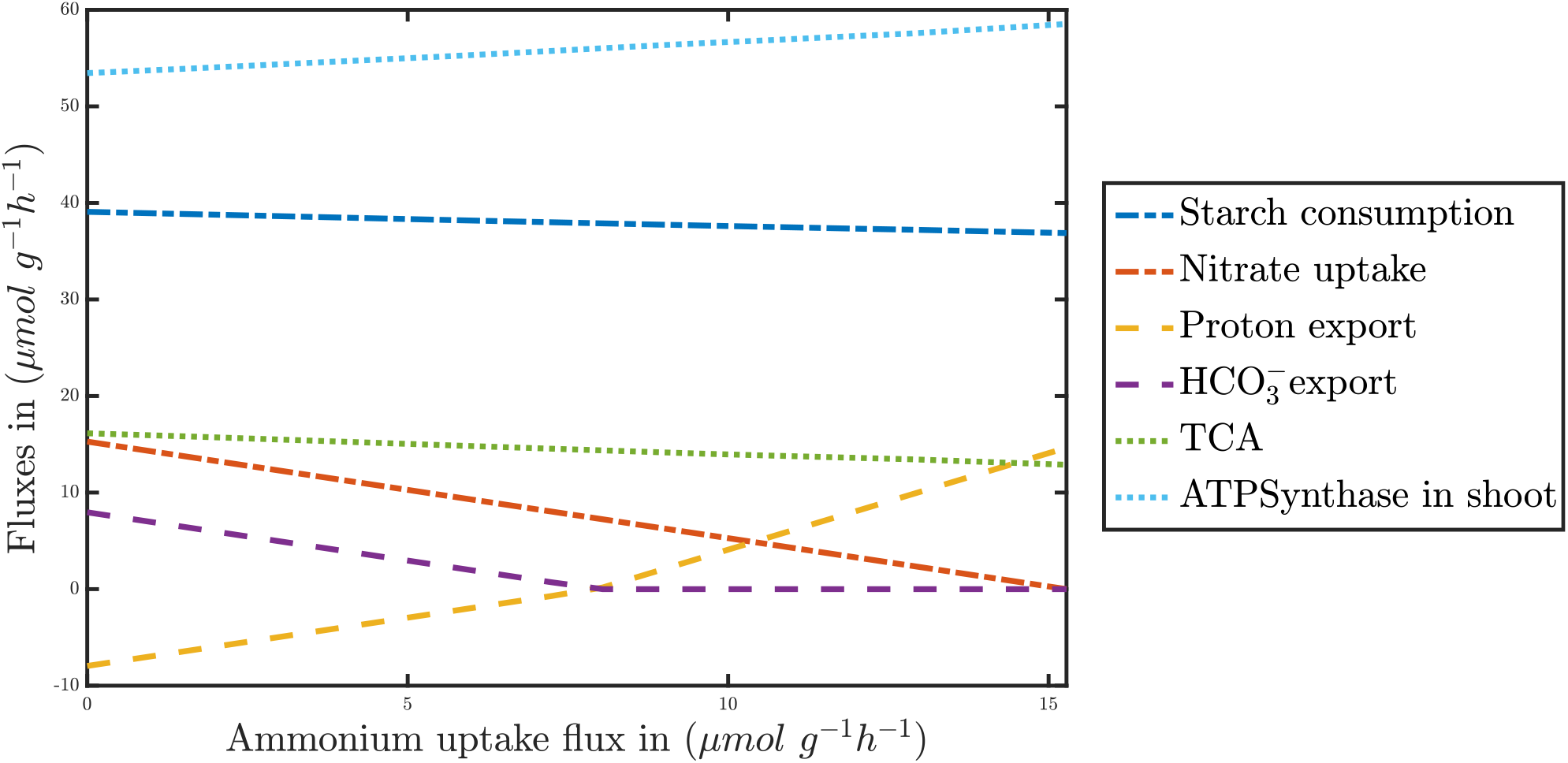
Responses of metabolic fluxes for increasing availability of ammonium. Maximal ammonium uptake (x-axis) was systematically varied and the response of selected fluxes depicted (y-axis). The fluxes were calculated by fixing the biomass production to 0.1 g/gDW per day and minimising the total quadratic flux through the network.

#### Symbiotic nitrogen fixation

We used the combined model (see the schematic in Figure 6) first to test under which conditions a symbiotic association with *S. meliloti* is beneficial to the host *M. truncatula*. For this, we compared the maximal predicted growth rate of the models with and without nitrogen fixing symbiont, for different external nitrate concentrations (see Figure 7). Clearly, the growth rate is slightly lower for the symbiotic system if sufficient nitrogen is available. This can be explained by the additional energy requirement to produce organic carbon to support the symbiont. However, under low nitrogen availability (in the form of nitrate or ammonium) the ability to fix nitrogen by the rhizobium allows for plant growth even without any external nitrogen source present. Thus, this combined model can explain and to some extent quantify under which conditions a symbiotic relationship is advantageous.

**Figure 6:**
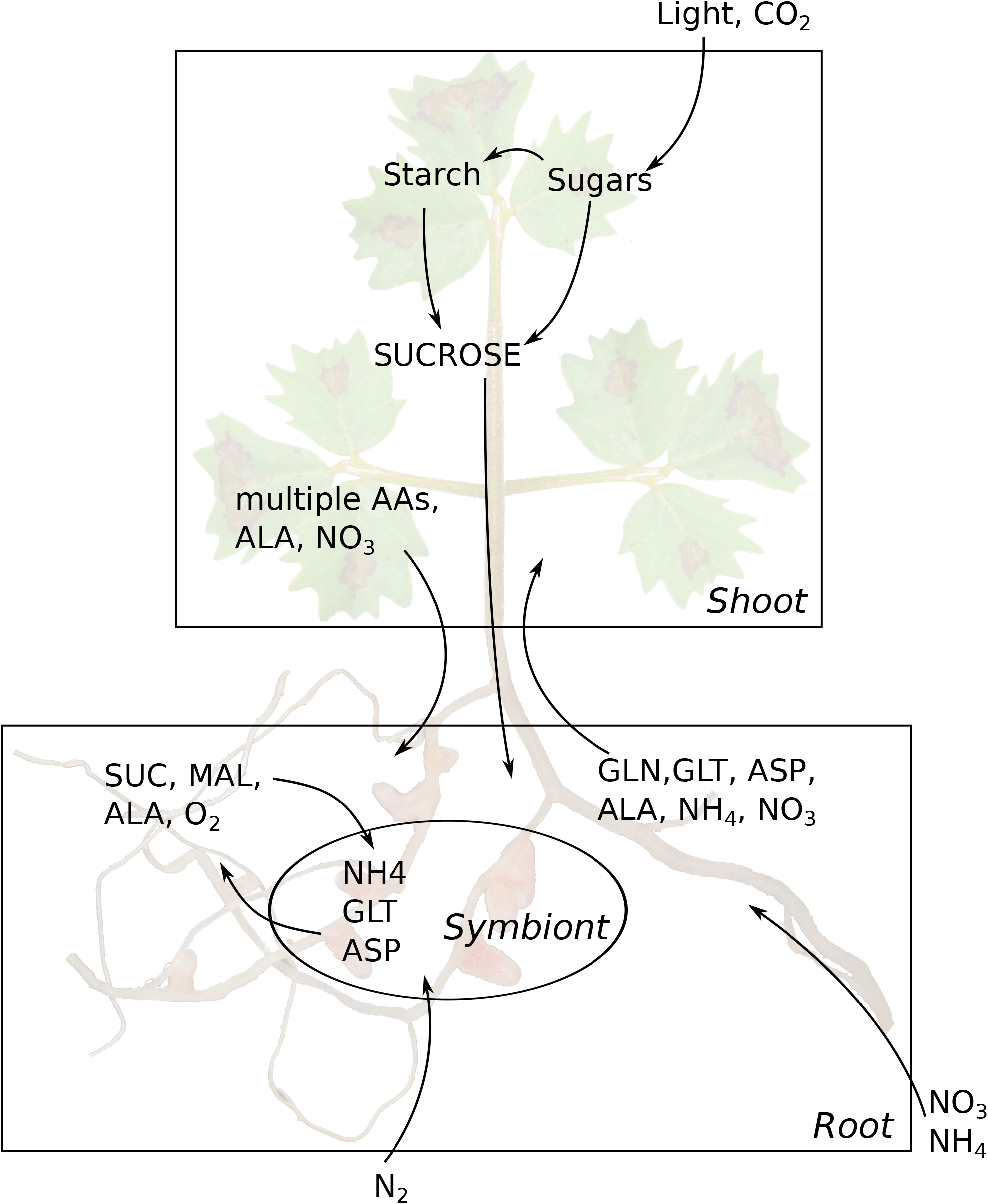
Layout of the two-tissue model combined with its symbiotic partner. The most important exchanges between the tissues and between plant and symbiont are shown.

**Figure 7:**
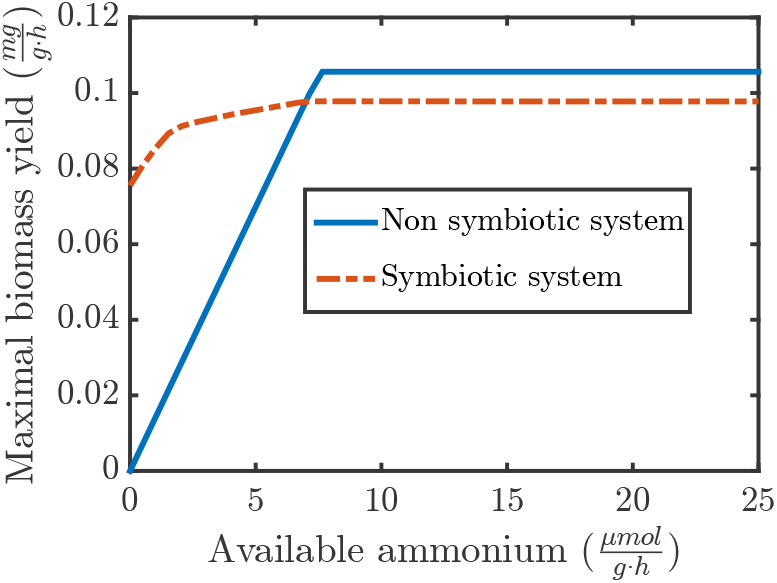
Comparison of the maximal growth of the symbiotic and non-symbiotic system. While the symbiotic system can grow without available ammonium by fixing nitrogen, this advantage necessitates a maintenance of the rhizobial symbiont, which reduces the energy available to the plant, leading to a lower maximal growth when sufficient ammonium is available.

There are multiple propositions in the literature concerning the transport of nitrogen from the rhizobium to the plant (Waters et al., 1998; Poole and Allaway, 2000; Lodwig et al., 2003). An initial investigation showed that the amount of oxygen available to the bacteroids is a limiting factor for the amount of nitrogen that can be fixed. However, while a higher oxygen concentration provides more energy to the bacteroid due to a higher activity of oxidative phosphorylation, the nitrogenase complex becomes irreversibly inactivated if oxygen concentrations are too high (Dingler and Oelze, 1985). While an increased potential biomass production is the most obvious effect of a better oxygen supply it is also interesting to investigate how the exchange fluxes between bacteroid and plant are affected if the amount of fixed nitrogen remains constant, but oxygen concentrations varies. For this, we fixed the growth rate of the plant to 80% of the maximal growth rate (determined above), and systematically varied the oxygen availability for the bacteroid and calculated the fluxes based on the assumption that enzymes are used in a maximally efficient way (see Methods).

Under a wide range of oxygen uptake rates, the model predicts extensive amino acid cycling (see Figure 8A). Alanine is predicted to be the only nitrogen-containing export product (for a schematic of the reactions in the symbiont see Figure 9). The amino group of glutamate is transferred by transaminases to pyruvate, yielding the export product alanine and ketoglutarate. The latter is fed into the TCA cycle, where one ATP is generated by succinyl-CoA synthase. Newly fixed nitrogen is exported as alanine, which is synthesised *de novo* from pyruvate by alanine dehydrogenase. Pyruvate in turn is produced by reverse action of pyruvate carboxylase. The standard Gibbs free energy of reaction of pyruvate carboxylase is *ΔG^0^* = — 1.37kcal/mol (according to MetaCyc), but due to the comparatively large amounts of dicarboxylic acids and the high demand for ATP it appears plausible that the equilibrium can easily be shifted to pyruvate. The imported carbon source required for pyruvate synthesis depends strongly on the available oxygen. For low oxygen supply (left region in Figure 8A), the capacity of the respiratory chain is limited and import of malate minimises the amount of produced NADH. With increasing oxygen availability, ATP production by oxidative phosphorylation is increasingly efficient. This explains the observed switch from malate to succinate uptake, because introduction of succinate into the TCA cycle provides one additional reductant as compared to malate (right region in Figure 8A). Therefore succinate is only used when sufficient oxygen is available (or when no malate is provided). In our simulation alanine was the only nitrogen export product from the rhizobia. However, there have been experiments in which alanine dehydrogenase was knocked out, showing that *de novo* alanine synthesis is not required for symbiotic growth (Allaway et al., 2000; Kumar et al., 2005). Our model allows a rapid reproduction of this experiment *in silico*. Figure 8B shows, even under these conditions, a substantial export of alanine, which is directly cycled against glutamate. However, in the knock-out simulation, nitrogen is exported from the symbiont in the form of ammonia, which is subsequently assimilated in the plant. In the knock-out conditions, the minimum amount of required oxygen increases, which corresponds to a slightly decreased growth, a phenomenon observed by Allaway et al. (2000) but not by Kumar et al. (2005). Interestingly, the knock-out simulation does not show any uptake of dicarboxylic acids at minimal oxygen concentrations but relies solely on the use of glutamate as carbon source. This can be explained by the fact that using glutamate as carbon source allows, as described above, the use of succinyl-CoA synthase to regenerate ATP, while simultaneously producing reductants. Thus, without the ability of a *de novo* alanine production, an optimal ratio of ATP to reductant is provided by pure glutamate uptake. With additional oxygen, the use of succinate as reductant donor and oxidative phosphorylation as ATP source becomes more efficient.

**Figure 8:**
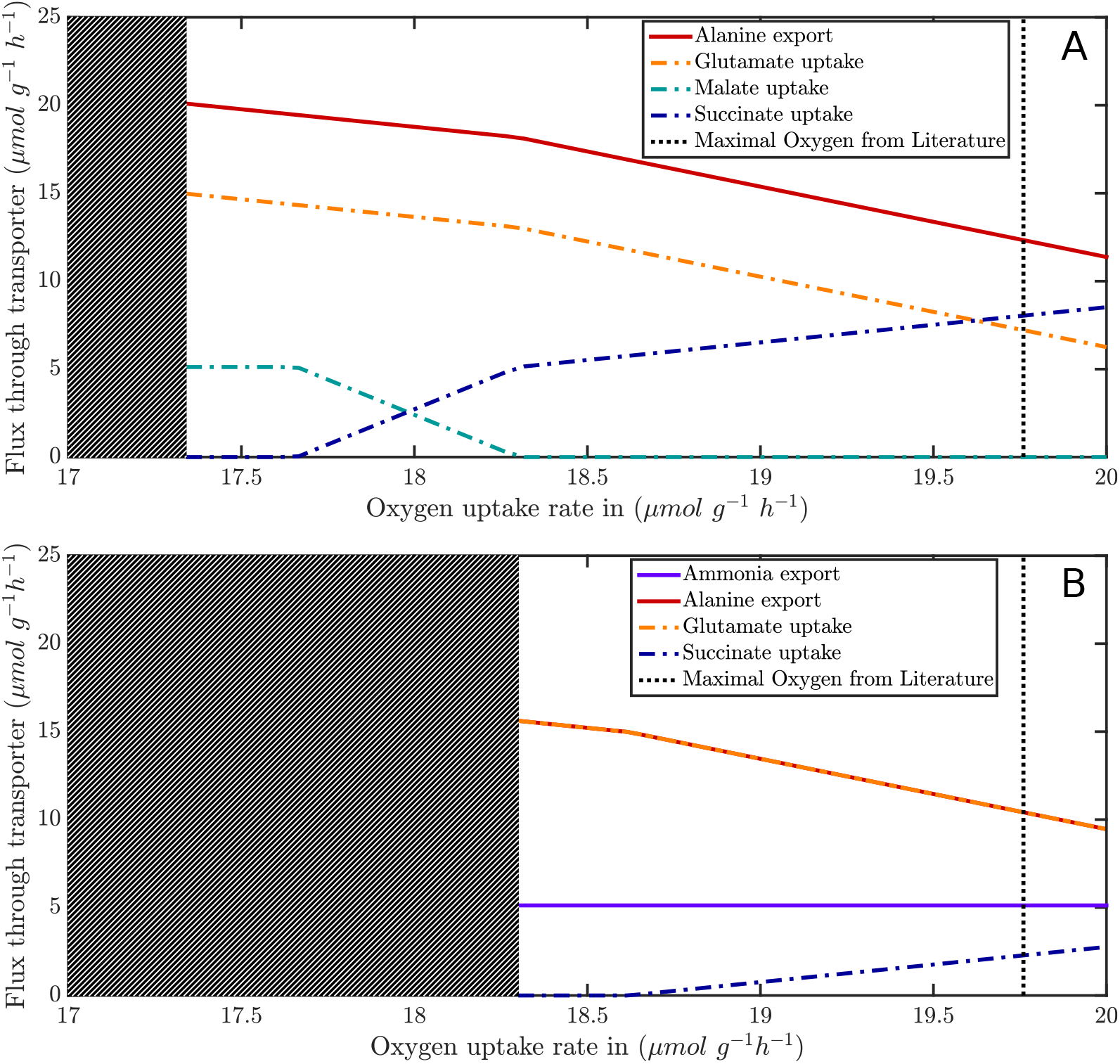
Effects of oxygen limitation on the exchange of compounds between plant and rhizobium. The hatched area indicates oxygen concentrations which do not allow sufficient nitrogen fixation for the assumed growth. (A) Scan with alanine dehydrogenase, in which alanine is the compound that exports the fixed nitrogen from the symbiont. (B) Scan without alanine dehydrogenase. While alanine is still exported (the same amount as glutamate uptake), ammonia is the exported product of nitrogen fixation.

**Figure 9:**
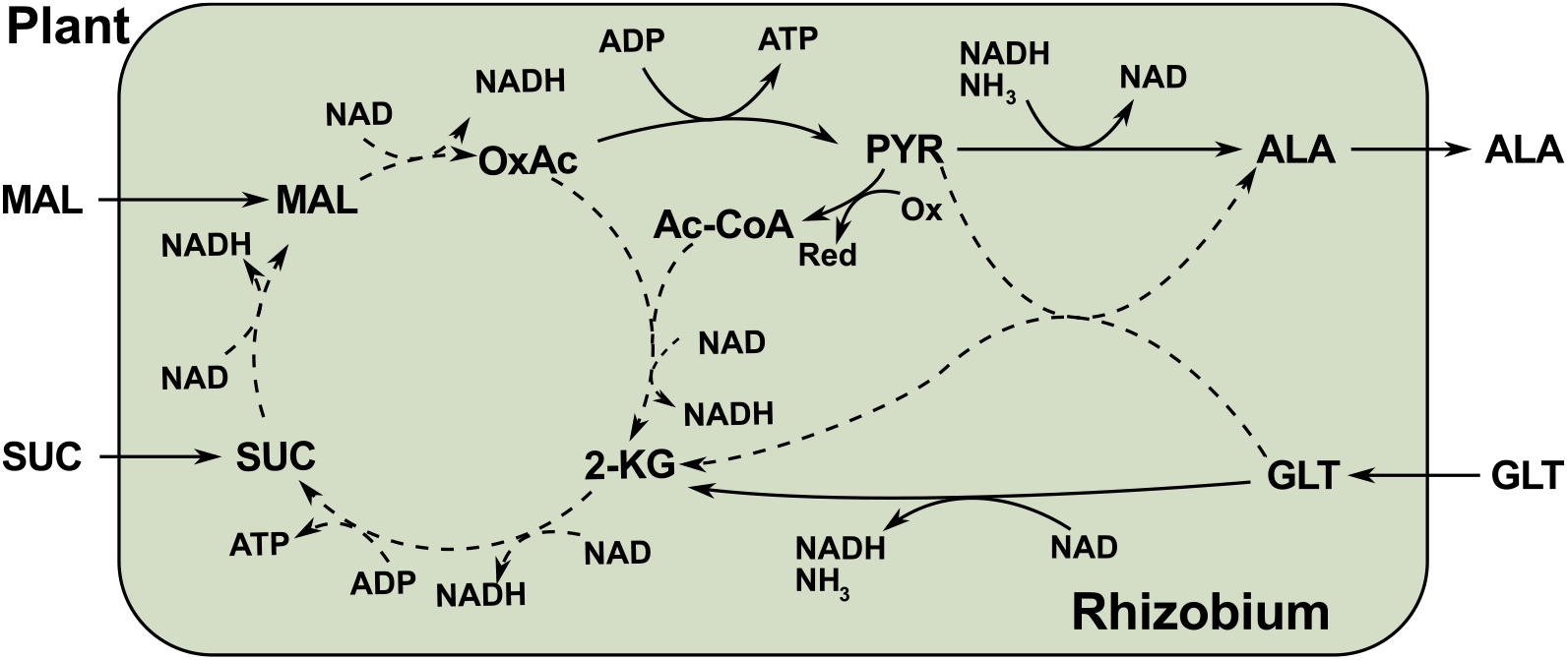
Overview of the main reactions occurring in the rhizobium. When glutamate is imported as carbon source, the TCA cycle is fed with ketoglutarate and one ATP is produced during conversion to succinate. The amino group of glutamate is directly transferred to alanine. If dicarboxylic acids are imported, less ATP is produced. Simultaneously, less reductant (mainly NADH) is produced, which is particularly pronounced for malate as carbon source. This reduces the flux through the respiratory chain, allowing the sybiont to better cope with limited oxygen.

## 3 Discussion

We have developed a genome scale compartmentalised model for the clover *Medicago truncatula*, a model plant for the legume–rhizobia symbiosis and connected this model to a rhizobial symbiont. We have carefully verified that our model is thermodynamically feasible, that all relevant intermediates can be replenished, and all reactions are balanced with respect to both mass and charge, which allows the interpretation of results regarding the effect of different types of nitrogen nutrition on the environmental pH. The reconstruction process was useful in its own right, as it resulted in the putative correction of a number of existing annotations. However, new annotations are not the primary objective for building detailed genome-scale metabolic models. More importantly, a genome-scale model provides a theoretical framework in which experimental observations can be interpreted and understood. Computational analyses allow to query the model, assess its capabilities, and enable novel interpretations of experimental observations in a theoretical context. An interesting observation is derived by our calculation of the theoretically maximal conversion rate of starch-derived carbon into biomass at night. Considering only energy and redox requirements for the formation of biomass, a remarkably high percentage of starch carbon (92% for growth on ammonium, 79% for growth on nitrate) can be converted into biomass. These numbers are of course overestimated and these conversion rates are not expected to be observed in nature. Still, these theoretical deliberations provide novel insight because they allow for a new interpretation of observed growth rates in the context of the largely unknown maintenance requirements. Experiments performed in *A. thaliana*, in which respiration and starch degradation rates were carefully measured throughout the night (Pyl et al., 2012), showed that at least 45% of the starch is respired. This value is still more than twice as high as the predicted minimum of 21% for the growth on nitrate. Moreover, Pyl et al. (2012) showed that the ratio of respired starch-derived carbon is highly dependent on the night temperature. The lowest value of 45% is observed for low (12°C) temperatures, while for nightly temperatures of 24° C (which was the same as the applied temperature during the day), the ratio of respired carbon increased to 75%. Estimating from our calculations that around 20% of carbon contained in starch need to be respired to build biomass, this allows to conclude that between 25% and 55% of starch-derived carbon is respired during the night for maintenance, depending on temperature and probably other external factors. This is in good agreement with previous findings of maintenance requirements of around 40% (Williams et al., 2010). The derived energy is required for processes, which are not directly related to growth and not included in our model, such as degradation and repolymerisation of proteins and mRNA, as well as transport processes across intracellular membranes or phosphorylation in regulatory pathways. But why the ratio is so strongly dependent on the ambient temperature remains unclear. To further understand the detailed requirements for maintenance energy, it will be necessary to formulate mathematical models of maintenance requirements based on experimental measurements of protein and mRNA turnover and intracellular transport (Piques et al., 2009; Stitt, 2013). The results of such models would allow to define further constraints on genome-scale metabolic models as the one presented here, and lead to a more profound understanding of the regulation of metabolic fluxes responding to environmental changes.

One future goal of our work is to explore the nature of the symbiosis between legumes and nitrogen fixing bacteria. As a first step, we therefore studied the responses of metabolic fluxes to changes in availability of nitrogen sources. If both nitrate and ammonium are abundant, the model predicts an exclusive uptake of ammonium, because integration into amino acids requires considerably less reductants and energy when compared to nitrate. This result is in agreement with the experimental observation that nitrate uptake is inhibited when ammonium is available (Ohmori et al., 1977). Thus, in an evolutionary context, our model suggests that the reason for this inhibition is an increased energetic efficiency. Simultaneously proton export is observed which is in accordance with soil acidification when ammonium fertilizers are used (Alam et al., 2007). If ammonium becomes limiting, the modelled fluxes change gradually to an increased uptake of nitrate, which is accompanied by a higher energy and reductant demand that is met by increased respiration during the night. The balancing of charges, using both ammonium and nitrate, might contribute to the observation that the presence of ammonium in many plants (de la Haba et al., 1990) inhibits nitrate uptake but does not turn it off completely, while from an energetic consideration, nitrate should not be used at all. At the same time, we have to be aware of the limitations of FBA when interpreting computational results simulating changes in environmental pH. In particular, ionic barriers commonly employed by plants cannot be modelled in FBA.

By combining the plant network with a rhizobial symbiont, we were able to illustrate the evolutionary advantages in undergoing a symbiosis and to simultaneously show that it can come at a cost when formed in a rich environment. We could demonstrate that the observations of amino acid cycling and alanine as nitrogen carrier fit into a paradigm of efficiency. Further, we were able to suggest that the reason for the use of alanine as export product lies in the ability to remove some surplus reductant from the bacteroid system, but that it is not strictly necessary for symbiotic growth. However, it is interesting that even when alanine dehydrogenase is knocked out, we observed the export of alanine (now exchanged with glutamate), which supports the hypothesis that alanine is a major export product in planta, even without being the nitrogen carrier.

## 4 Materials and Methods

### 4.1 Metabolic Model Construction

The model construction procedure is summarised in Figure 10. Depicted are key steps in model development and the required data source to perform each step.

**Figure 10:**
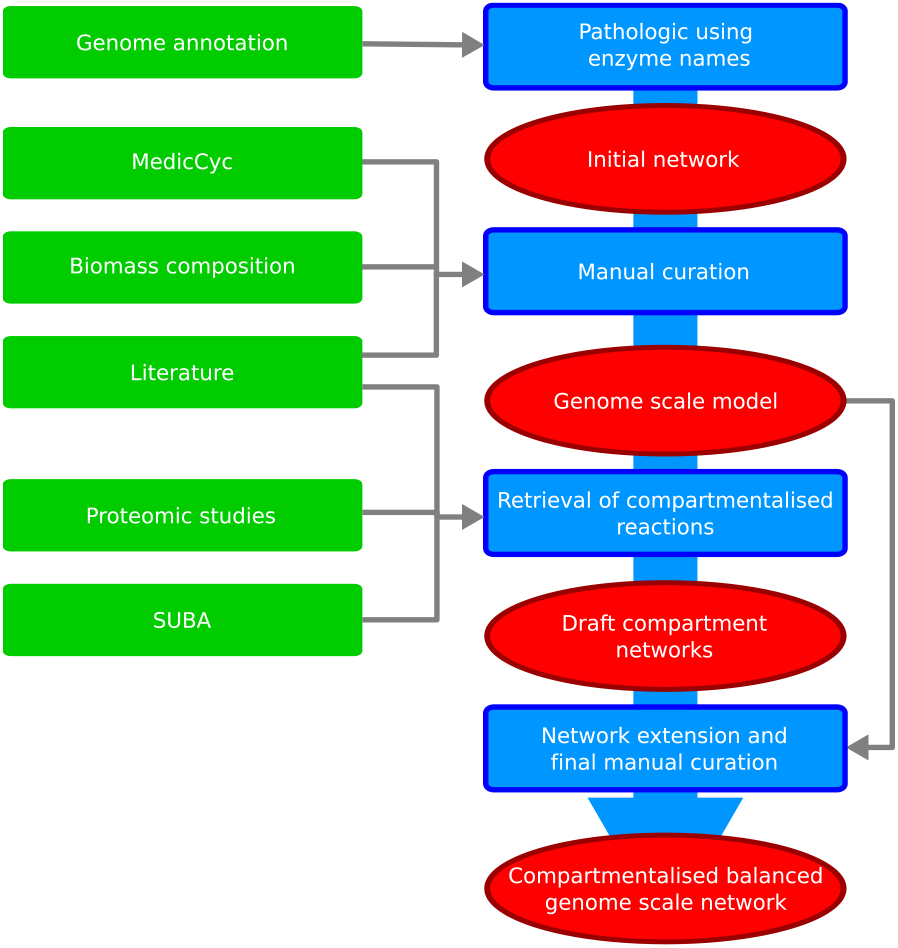
Layout of the model generation process. Data sources are marked in green, model stages in red and process steps blue. During manual curation producibility of all measured compounds was ensured and all reactions where charge balanced.

#### Initial Network Generation

The model is based on version 3.5v5 of the *Medicago truncatula* genome annotation (Young et al., 2011). The initial construction was performed using PathoLogic enzyme name matching from the PathwayTools program (Karp et al., 2010) and makes use of the MedicCyc (v1.0) database (Urbanczyk-Wochniak and Sumner, 2007), which is based on the BioCyc (Caspi et al., 2010) suite of databases. Information from MedicCyc (v1.0) was incoporated based on homology analysis between gene information in MedicCyc and the current genome annotation. This resulted in a list of enzyme-catalysed reactions converting mainly small molecules. However, the MetaCyc database also includes a number of reactions which contain macromolecules. In general, metabolic network models are limited to reactions acting on small molecules. Macromolecules, such as proteins or glucans, are not specifically included because of the enormous number of theoretically possible polymers. However, some macromolecules play important roles in metabolism and require special treatment. Among these are in particular small carrier proteins such as ferredoxins (for charge transfer reactions) or acetyl carrier proteins (for fatty acid biosynthesis). As essential cofactors, these molecules have to be explicitly included in the reaction stoichiometries but their synthesis pathways are outside the scope of a metabolic network model. To verify mass balance of reactions involving such molecules, we exploit that these macromolecules in their various modified forms form conserved moieties, i. e. that their core structures are not changed by any reaction in the network. For reactions involved in synthesis or degradation of polymeric sugars, such as starch, we replaced the compounds representing the polymers of a non-specified degree of polymerisation (such as starch(n)) by the monomers in such a way that the overall reaction is correctly balanced. To this extent, a xylan polymer for example is assumed to consist of dehydrated xylose subunits (C5H8O4).

Some reactions added to the model by PathwayTools had poor evidence. The name matching algorithm adds reactions if the assigned enzyme names in MetaCyc match those in the genome annotation, which leads to problems with unspecific annotations such as “alcohol dehydrogenase” or “aldehyde dehydrogenase”. Such descriptors resulted in the addition of many reactions that were disconnected from the remaining network. We therefore removed reactions from the model which fulfilled all of the following conditions: 1) a contained metabolite is only present in this reaction, 2) the reaction cannot carry any flux under the condition that all nutrients can be imported and all metabolites can be exported, and 3) there exists no evidence for the presence of the reaction beyond a match of unspecific reaction names.

#### Gap filling

Gaps in the initial network reconstruction were identified by verifying that all experimentally observed metabolites were actually producible from the nutrients. For this, we collected a list of experimentally verified metabolites from various studies (Barsch et al., 2006; Broeckling et al., 2005; Farag et al., 2007; Huhman and Sumner, 2002; Kowalska et al., 2007; Pollier et al., 2011). We then computed for each of these metabolites whether a stationary flux distribution exists (see below) such that this metabolite is produced while only the nutrients are consumed. When this check failed, the MetaCyc database was searched for plausible candidate reactions which needed to be added in order to allow for a production of the respective metabolite. The identified reactions were manually added to the network.

#### Atomic and Charge balancing

We intended to create a model which can simulate both photosynthetic and heterotrophic growth. Because proton gradients are essential in both regimes, we needed to ensure all reactions are charge balanced. Using MetaCyc charge information (all compounds protonated to a pH of 7.3) we checked all reactions for their charge balance. Apparently, several MetaCyc reactions have been curated to achieve mass balance by adding protons. This is in particular true for reactions in the phospholipid desaturation pathway, but also in reactions representing dehydrogenase or mixed function oxygenase activities. Where possible, we curated these reactions by assuming common desaturase, dehydrogenase or mixed function oxidase activities. In general, when protons had been added for mass balance reasons they were replaced by NAD(P)H + H+/NAD(P)+ pairs to achieve charge balance. This replacement was done by applying the general rules which are laid out in Table III.

**Table III:**
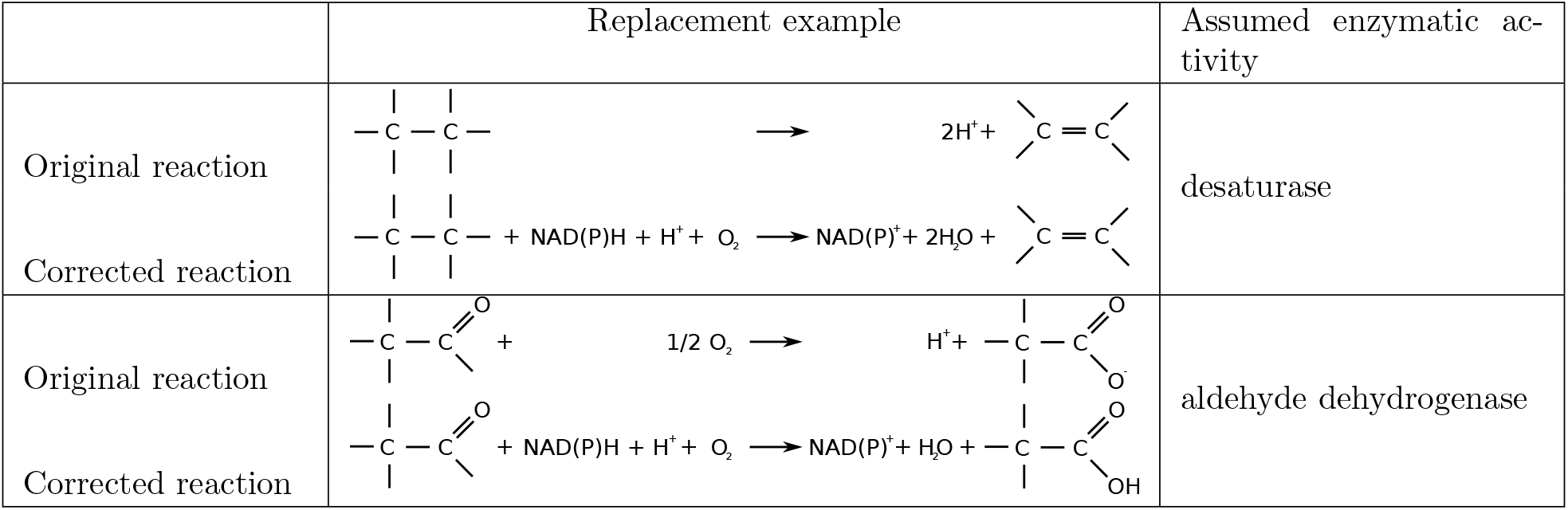
Examples for the replacement of reactions that are not charge balanced by plausible enzymatic activities, which ensure a correct charge balance.

Atomic balance was checked using the method detailed in Gevorgyan et al. (2008) and only light was shown as unconserved. This is to be expected, as light does only transfer energy into the system and does not contribute towards atomic balance.

### 4.2 Model Compartmentalisation

In order to realistically describe the metabolism of a plant cell, it is important to develop a fully compartmentalised model. Each compartment fulfils specialised tasks while key processes, such as ATP generation, are only possible due to gradients established over inter-compartmental membranes. Key challenges when building a compartmentalised model are a) to realistically assign compartments to each reaction, b) to define transport reactions over intracellular membranes and c) to ensure that the resulting model is not able to generate and maintain gradients which are thermodynamically infeasible.

#### Assigning compartments to reactions

To overcome the first challenge, we applied the following approach: We first scanned all available protein sequences for annotated genes against the *Arabidopsis thaliana* genome on TAIR (Lamesch et al., 2012) and extracted information about localisation from the SUBA database (Heazlewood et al., 2007). We assigned compartments to only those reactions for which experimental evidence, such as GFP-localisation, MS identification, TAIR data, and swissprot data, existed. Because computational predictions are known to result in a large number of false positives, they were not included during this step. In addition, two recent proteomic studies for root plastids (Daher et al., 2010) and mitochondria (Dubinin et al., 2011) in *M. truncatula* were used to assign compartments to reactions. For this, the provided gene identifiers, which were based on different versions of the Medicago truncatula Gene Index (Quackenbush et al., 2001), were translated to gene identifiers of version 3.5v5 of the *M. truncatula* genome annotation using BLAST. All these sources of information were integrated and if evidence for the presence of a protein in a specific compartment was found in at least one source it was added to this compartment. This initial process resulted in a draft compartmentalised network with still a large number of reactions not yet assigned to compartments. In a next step, we used an established network modelling approach (Christian et al., 2009) to ensure that each compartment contains a self-consistent metabolic network: The network extension developed by Christian et al. identifies a minimal set of reactions from a reference network which need to be added to a draft network in order to ensure that a number of ‘seed’ metabolites can be metabolised into a number of ‘targets’. While for a whole organism, the seed can directly be inferred from defined growth media, and the targets can be obtained by metabolomic data, their definition is not straight-forward for single compartments. We obtained seed and target sets for each compartment by an extensive literature research for metabolites which are known to be imported into the compartment or produced by the compartment. This step was performed for all compartments except the cytosol and the vacuole. This procedure is difficult for the cytosol, because it is in direct contact to all other compartments. Similarly, the vacuole is a main location for degradation and storage, making a meaningful definition of seeds and targets difficult. For the vacuole we therefore used the gene localised reactions for the initial compartment. We manually assigned additional reactions only if they were missing steps in degradation pathways for which a majority of reactions was already present in the vacuole. Whereas in network extension for whole organism networks, usually a network containing all reactions stored in a biochemical database is used as a reference, we here use all reactions of the uncompartmentalised network as reference, because our goal is to assign each reaction a particular compartment. The lists of seed and target metabolites used for this process and the resulting reactions added to each compartment can be found in Supplementary Material 1. All reactions which remained without assigned compartment after applying network extension to all compartments were subsequently assigned to the cytosol, with the exception of the sterol metabolism which was assigned to the ER.

#### Transporters

Describing transport processes of metabolites over membranes is a challenge in all compartmentalised metabolic networks. The main difficulty is that exact knowledge about transported metabolites is still missing. Further, the existence of several transporters has to be assumed without experimental evidence due to localisation of biosynthetic enzymes. Recently, Linka and Weber reviewed many characterised transport systems for plants (Linka and Weber, 2010). However, even this comprehensive review must necessarily miss transporters for which no molecular characterisation exists yet, but which are essential for production of different metabolites. We added transporters between compartments to ensure that all metabolites which were producible in the uncompartmentalised network were still producible in the compartmentalised network. For this, we used the information given in Linka and Weber (2010) complemented by several other sources (Babujee et al., 2010; Helliwell et al., 2001; León and Sánchez-Serrano, 1999), and otherwise included transporters which seemed biologically most plausible. In the presented model, we assume that most transporters do not carry a net charge. This is true in many instances, but more detailed information on the charges carried by transporters would further improve the model description and provide further insights (Cheung et al., 2013). A full list of added transporters is given in Supplementary Material 5.

#### Maintaining thermodynamic integrity

Another challenge arises with protonation states of different compounds. This is particularly important for the mitochondrial compartment which harbours the electron transfer chain for ATP synthesis. Because there are plenty of transporters over the mitochondrial membrane, we had to ensure in the model description that these transporters do not allow a net-charge exchange over the membrane, because otherwise proton gradients might be generated within the model by a simple shuttling of metabolites. This would result in the generation of ATP uncoupled from energy provided by exergonic reactions, thus rendering the model thermodynamically infeasible and predictions regarding ATP generating fluxes meaningless. All transporters over the mitochondrial and plastidial membranes were therefore balanced for charges by adding protons in a way that a net charge of zero is translocated. The only exception to this were reactions from the electron transfer chain for which proton translocation is known. Similarly, import and export of metabolites over the cellular membrane is critical for charge balancing the system. We include uptake and export of charged particles by importers and exporters. Additionally, we introduce a proton and bicarbonate exchanger. In our simulations, we applied the constraint (see below) that the net charge exchange is zero. This resulted in the model behaviour that protons and bicarbonate were used to balance excess uptake or export of charges. The biomass reaction itself was charge-balanced by adding protons to ensure that the net charge of the biomass is zero.

### 4.3 Generation of a two tissue shoot-root Network

To more realistically describe processes taking place in root cells, we have constructed a tissue-specific subnetwork. We used the presence calls provided in Benedito et al. (2008) to determine core reactions sets for shoots and roots. We assume a gene to be present if it is determined as present in a majority of experiments. Shoot core genes are the combination of all genes present in stems and leaves. Root core genes are those genes determined as present in the root. The core reaction sets used for the model generation were determined by matching the determined genes using the gene-protein-reaction association rules in the model, and determining which reactions could be activated by the present genes. The model was then duplicated as a root and a shoot model and exchangers between the two tissues were added similar to the concept detailed in Gomes De Oliveira Dal’molin et al. (2015). A full list of tissue exchanges is provided in the supplemental files. All uptakes from the external medium except for CO_2_, light and starch were removed from the leave model, and glucose, starch, light and sucrose exchangers were removed from the root model. This combined model was made consistent, i.e. all reactions which could not carry flux were removed using FASTCC Vlassis et al. (2014). The consistent model was then subjected to the FASTCORE algorithm (Vlassis et al., 2014). FASTCORE extracts a small consistent network from the original network, in which all reactions from a given core set can be activated. We used those reactions as core for which the models gene-protein-reaction relationships returned true, if the core genes for the respective tissue were set as active. In addition, some reactions were turned off to generate the individual models. This step was repeated four times with slightly modified models to obtain four individual conditions:

1. Growth using ammonium as sole nitrogen source during the day.
2. Growth using ammonium as sole nitrogen source during the night.
3. Growth using nitrate as sole nitrogen source during the day.
4. Growth using nitrate as sole nitrogen source during the night.

For day conditions, the biomass reaction without starch in the leaves, and starch import were turned off, while night conditions had the starch containing biomass and light import turned off. For growth on nitrate, the ammonium uptake was shut down and for growth on ammonium, the nitrate exchanger was set to 0, respectively. Day conditions implied production of a biomass containing starch along with availability of light, while night conditions only allowed starch as a carbon and energy source while producing a biomass without starch. Finally, all reactions present in at least one of these four models were combined to yield the two tissue model. Performing only one FASTCORE calculation could have led to disconnected subnetworks, where e.g. light could be used to fix carbon but the carbon might not be linked to the remaining network. We employed multiple constraints for the analysis of the model. An important factor, atp maintennce, was estimated based on data from De Vries (1975) for *Trifolium repens*, white clover, the closest relative for which data is available. This plant needs 15 mg glucose per gram dry weight per day for maintenance (De Vries, 1975). With a regenerating ability of about 32mol ATP per mol glucose, this translates to an hourly maintenance cost of about 111.1 μmol/(gh), a number which has to be distributed between the two tissues. With an approximate 1:2 ratio (root:shoot) the atp maintenance for shoot was set to 74μmol/(gh) and the root maintenance set to 37μmol/(gh) Similarily the biomass production combines two thirds of a shoot biomass and one third of a root biomass to one μmol of biomass, with a weight of 1g. We assumed a normal growth rate of 0.1g/(gd) based on data for *M. sativa* (Lötscher et al., 2004) for simulations where a fixed biomass production was required. The maximal starch consumption in leaves was set to 46.32 μmol/(gh), based on the starch storage in our biomass experiments during the day. The maximal rate of photon uptake was set to 1000μmol/(gh).

### 4.4 Building a symbiotic system

A network for *Sinorhizobium meliloti* 1021 was extracted from MetaCyc. The model was curated to be able to produce most biomass components from an *E.coli* biomass defintion (Orth et al., 2011). The model was connected to the plant root submodel using exchange reactions present in the literature (for Reviews see e.g. Udvardi and Day (1997); Prell and Poole (2006); Udvardi and Poole (2013) and a list of used transporters can be found in the supplemental data).

To be able to compare fluxes in the rhizobium with fluxes in the plant, we had to adjust all fluxes to use a common unit, which we based on plant dry weight. To achieve this we used the following data: The ratio between nodule dry weight and plant dry weight is in the range between 0.01 g/g (Vance et al., 1979) and 0.035g/g (Aydi et al., 2004). In addition, there are roughly 8 o 10^10^ bacteroids per gram nodule (Sutton et al., 1977) and a bacteroid weights approximately 3·10^−12^ g (Bergersen, 1997). Thus the total amount of bacteroids per gram of plant dry weight is in between 2.40 mg/g and 8·40 mg/g. Table I lists the original and adapted ranges of fluxes. We assumed a symbiont maintenance energy of 7.6mmol/(gh) based on symbiont weight which translates to 63.47μmol/(gh) based on plant weight. An overview of the model including the symbiont compartment and the main exchangers is shown in Figure 6. This is the maintenance energy commonly used in *E.coli* (Orth et al., 2011). After combining the models, we noticed, that our biomass formulation has a very large asparagine fraction. This is likely due to the fact, that we fed our plants with excessive amounts of nitrogen to avoid any nodulation. Therefore, we reduced the amount of free asparagine in the plant biomass formulation to 5% of the original amount. Without this adjustment, the maximal growth rate with maintenance energy is at about 0.04g/(gd), which is about 53% lower than with the adjusted amount. In addition, the rhizobium spotted a citrate lyase reaction, which by acting in reverse allowed the production of some ATP. Since this protein commonly catalyses the forward reaction, we set its lower bound to zero. For our simulations, the symbiont was assumed to have finally differentiated and therefore stopped growth, i.e. not consume the nitrogen it fixes, de facto becoming an additional compartment of the plant.

### 4.5 Network analysis

Genome-scale networks are most conveniently described by the stoichiometry matrix N, in which columns represent biochemical reactions and rows correspond to metabolites, and the coefficients define whether a metabolite is consumed (negative) or produced (positive) by a particular reaction. The dynamic behaviour of the metabolic network is governed through the stoichiometry matrix by

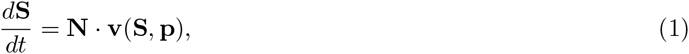

where **S** is the vector of metabolite concentrations, **v** is the vector containing all reaction rates and **p** is a vector with all kinetic parameters. Assuming that the system is in stationary state, the dynamic equation is reduced to the stoichiometric equation

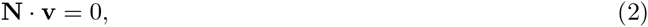

which allows to derive statements about the flux distribution v from the stoichiometry matrix alone. This system is usually under-determined, meaning that many flux distributions fulfil the equation. To restrict the solution space, further constraints are defined. These reflect thermodynamic constraints restricting the direction of some reactions, as well as experimentally observed constraints, such as maximal uptake rates and measured growth rates combined with biomass composition. Such limitations are in the most general form described by inequalities

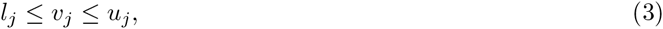

where *l_j_* and *u_j_* are lower and upper bounds restricting the flux through a reaction or transporter *j*. In our case, an important constraint also reflects the fact that the net charge accumulation must be zero. This constraint is written as

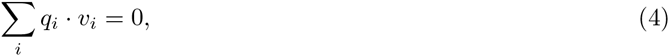

where *q_i_* is the net charge translocated over the cellular membrane by transporter *i*. Network analysis was performed using ScrumPy (Poolman, 2006) (mudshark.brookes.ac.uk/ScrumPy) and the COBRA tool-box (Schellenberger et al., 2011) with CPLEX as solver. A ScrumPy to CPLEX interface is provided in the supplements.

#### Producibility of metabolites

To verify whether a particular metabolite X is producible, a reaction 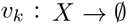 is added to the network. Then, it is tested whether a solution vector v exists, such that the stationarity condition 2 and the constraints 3 and 4 are fulfilled, for which *υ_k_* > 0. If the system is solvable, there exists a stationary flux distribution which consumes only nutrients and produces (at least) metabolite *X*.

#### Energy and redox requirements

To find a theoretical minimum of energy and reductant requirements to produce a certain metabolite k, we introduce as above a reaction 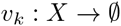 and two reactions consuming (positive flux) or producing (negative flux) energy and redox equivalents (the ATPase, ATP + H_2_O + H^+^ → ADP + Pi, and the NADPH dehydrogenase, NADPH + H^+^ + 0.5 O_2_ → NADP^+^ + H_2_O). The optimisation problem is to find a flux distribution which produces one unit flux through the reaction consuming metabolite *k*, while minimising the fluxes through the ATPase and NADPH dehydrogenase. For this, a three step optimisation is performed. First both ATPase and dehydrogenase remain unconstrained with the optimisation being a minimisation of starch consumption. The next step is to fix starch consumption and minimise the flux through the dehydrogenase reaction. The final step is to also fix the dehydrogenase flux and minimise the flux through the ATPase reaction.

#### Flux Balance Analysis

The identification of special solutions, which optimise some plausible objective function, is subject of Flux Balance Analysis (FBA, (Kauffman et al., 2003)). The choice of the objective function is an unsolved problem. A common assumption is that fluxes are arranged such that biomass yield is maximised (Feist and Palsson, 2010). However, this assumption is experimentally supported only for *E. coli* (Edwards and Palsson, 2000; Edwards et al., 2001; Feist et al., 2007) and the objective is certainly debatable for eukaryotic and multicellular organisms. Different tissues of a complex organism and likewise different compartments in a eukaryotic cell will have different functions and therefore follow different objectives. Unless otherwise stated, we perform our calculations by solving the optimisation problem to find a flux distribution, that minimises the quadratic fluxes. Due to the properties of the quadratic problem, this solution is necessarily unique thus avoiding problems with alternate optima. While this objective may not reflect the true objective, it reflects a plausible assumption and allows to investigate how the most economic flux distribution changes upon external perturbations. To calculate theoretically most efficient conversions during night, we introduce ‘carbon optimisation’. Here, a flux distribution is determined which minimises export of carbon (e. g. as CO_2_) and thus ensures a maximal ratio of carbon from starch being incorporated into the biomass. In all simulations where flux minimisation was desired, the glucose exporter was set to zero, as otherwise, glucose is exported when starch is degraded. The unrealistic solution is obtained because if maltose is split into glucose and glucose phosphate and unphosphorylated glucose is exported, a slightly lower overall flux is obtained than for the more realistic solution in which the extra ATP is produced which is required to phosphorylate the second glucose.

#### Performance scans of metabolism

Some demands on the metabolic network are unknown. It is, for example, unknown how much energy is required for maintenance or how many reductants are required to maintain the redox balance. By fixing fluxes representing these requirements to a certain value and systematically varying its quantity we can investigate how the system responds to external challenges. For this, an additional constraint, *υ_i_* = *α*, is introduced and the equation system is solved for a certain range of values of *α*. To determine varying energy and redox demands, we introduce two additional reactions, an ATP hydrolysis (ATPase: ATP + H_2_O + H^+^ → ADP + Pi) and a dehydrogenase-like reaction (DEHOG: NADPH + H^+^ + 0.5 O_2_ → NADP + + H_2_O). When not mentioned otherwise, these reactions are The values of these fluxes are fixed and the system is solved for a range of values. Such scans underly Figs. 5 and 8.

### 4.6 Biomass experiments

#### Hydroponic culture of *M. truncatula*

Seeds were scarified for 8 minutes using 95/97% H_2_SO_4_. After washing they were incubated in a hydrochloric solution (Millipore tablets 2WCL01F50) for 30 minutes and finally washed with a large surplus of sterilized dH_2_O. The seedlings were transferred to agar plates (0.8% (w/v)) and left in the dark at 25 °C for 48 hours. Plants were grown hydroponically for eight weeks at 21 °C and 16/8h Light/dark cycle. The nutrient solution was prepared according to the *Medicago truncatula* Handbook (Barker et al., 2006) with a final ammonium nitrate concentration of 8 mmol L^−1^ and no potassium nitrate. The tubs were aerated with air stones. 8 week old plants were harvested in 2 intervals. One harvest at the end of the light period, the other at the end of the dark period. After harvest, plants were submerged in liquid nitrogen and freeze dried till further use.

#### Extraction of soluble metabolites

Soluble metabolites were extracted from lyophilized tissue using methanol/chloroform according to the protocol of Lisec et al. (2006). Tissue samples (leaves, root, stem) were transferred to a mortar and ground to a fine powder under liquid nitrogen. A known mass (25 mg) was weighed into a pre-cooled 15 mL Falcon tube and 7mL of 100% Methanol (pre cooled at −20 °C) was added and vortexed for 10s. 300μL of ribitol (0.2mg mL^−1^) was added as internal standard at this point. Tubes were shaken for 10 min at 70 ^°^C. 730 μL were transferred to 2 mL eppendorf tubes and the tubes were centrifuged at 11,000 g for 10min to pellet insoluble material. 375 μL chloroform and 750 μL water (precooled to −20 and 4 °C respectively) were added. The tubes were briefly vortexed and centrifuged for 15min at 2,200 g to allow the non-polar and polar phases to separate. For GC-MS analysis aliquots of the polar phase were taken and dried down. Lipids were extracted using Hexane/Isopropanol extraction according to Hara and Radin (1978) and quantified by weight.

#### Protein extraction, precipitation and hydrolysis

Protein extraction, precipitation and hydrolysis was performed according to Williams et al. (2008) using a Urea/Thiourea extraction. Analysis was performed by GC-MS. twice with 4 mL of ice-cold acetone and again centrifuged as above. The pellets were allowed to stand in a flow hood to evaporate any acetone left. Protein weights were taken after drying overnight. The protein pellets were re-suspended in 2 mL of 6M HCl and incubated in a heat block at 100 ^°^C for 24 hours for hydrolysis. Samples were cooled to room temperature and an aliquot (20 μL to 50 μL) of the hydrolysates (containing amino acids) were transferred to eppendorf tubes and dried in a vacuum dryer overnight to remove HCl. The dried samples were subjected to GC-MS analysis.

#### Nucleic acids

RNA and DNA were extracted using TRIzol/DNAse and Chloroform/isoamylalcohol extraction methods respectively (adapted from Sambrook and Russel (2000)). Quantification was performed photospectrometrically using a nanodrop (ND-1000 Spectrophotometer; NanoDrop Technologies Inc.; Wilmington, DE, USA).

#### Starch

Starch was extracted according to an established protocol (Williams et al., 2008). The insoluble pellet obtained after the methanol chloroform extract was used as starting point. Each sample was gelatinized by autoclaving the residue in 3mL of 25 mM sodium acetate for 3 h at 121 °C, 1.04 bars pressure followed by enzymatic digestion with 20 U α-amylase (Sigma-Aldrich) and 5 U amyloglucosidase (Sigma-Aldrich) for 16 h at 37 ^°^C. The samples were centrifuged and the supernatant containing the glucose monomers was collected, freeze dried and stored at −80°C until further analysis. Starch amounts were determined with an enzymatic assay.

#### Gas chromatography-mass spectrometry

GC-MS analysis was done as in Masakapalli et al. (2013) and Williams et al. (2008) using an Agilent 79890 GC coupled to an Agilent 5975 quadrupole MS detector, electron impact ionisation (70 eV) equipped with an Agilent HP 5-ms column (30 mm, 0.25 mm inner diameter) at the facility in the Department of Plant Sciences, University of Oxford, UK.

The protein hydrolysate extracts were derivatised by TBDMS (Antoniewicz et al., 2007) whereas the soluble extracts were derivatised by MeOX TMS (Lisec et al., 2006). To obtain the TBDMS derivatives, the dried samples were first dissolved in 25 μL of pyridine and incubated at 37 °C, shaking at 900 rpm for 30 min. Then 35μL of MtBSTFA + 1% t-BDMCS (N-methyl-N-(t-butyldimethylsilyl) trifluoroacetamide + 1% t-butyl-dimethylchlorosilane, Regis Technologies Inc) was added and the mixture was incubated at 60 ^°^C, shaking at 900 rpm for 30 min. To obtain the MeOX TMS derivatives, 40 μL of 20mg mL^−1^ methoxyamine hydrochloride (Sigma) in pyridine was added to the sample and incubated at 37 °C, shaking at 900 rpm for 2 h. Then 70 μL MSTFA (N-methyl-N-(trimethylsilyl)trifluroacetamide (HiChrom)) was added and the mixture was incubated at 37 °C for 30 min at 900 rpm. The derivatised samples were transferred to 8 mm glass vials (Chromacol) and sealed with a septum cap. The samples were run through GC-MS. The mass spectra of all the samples were acquired for m/z 146-600 by scanning (at 4.38 or 3.19 scans s^−1^). A solvent delay of 10min was set so that the mass spectrometer was turned on after elution of the bulk of the solvent from the column. analysed using GC-MS (derivatisation agent: MtBSTFA + 1% t-BDMCS). Lipids were extracted using a standard isopropanol/hexane extraction and quantified by weight. RNA and DNA were extracted using TRIzol/DNAse and Chloroform/isoamylalcohol extraction methods respectively. Starch was determined with an enzymatic assay (digestion followed by spectrometric assay).

## Supplemental Material

**Supplementary Information 1**: A list of seeds and target metabolites used in the network extension algorithm and the suggested extension.

**Supplementary Information 2**: Energy costs for all metabolites with exporters in the model.

**Supplementary Information 3**: Exchanged metabolites between shoot and root and symbiont and root.

**Supplementary Information 4**: A complete list of all single metabolite transporters in the model.

**Supplementary Information 5**: A zip file containing MATLAB scripts and the CPLEX interface for ScrumPy.

**Supplementary Information 6**: An sbml file containing the created metabolic network of *Medicago truncatula*.

## Author Contributions

OE,MP and TP conceived the study. NC and TP worked on the network compartmentalisation. TP and SM performed the biomass experiments. TP and OE performed the computational analysis and network reconstruction. TP, OE, NC and LS, wrote the manuscript. All authors edited the manuscript.

